# Proton-control of transitions in an amino-acid transporter

**DOI:** 10.1101/606913

**Authors:** Z. Wu, I. Alibay, S. Newstead, P. C. Biggin

**Affiliations:** University of Oxford

**Keywords:** Molecular dynamics, free energy, conformational change

## Abstract

Amino acid transport into the cell is often coupled to the proton electrochemical gradient, as found in the solute carrier (SLC) 36 family of proton coupled amino acid transporters (PATs). Although no structure of a human PAT exists, the crystal structure of a related homolog, GkApcT, from bacteria has recently been solved in an inward occluded state and allows an opportunity to examine how protons are coupled to amino acid transport. Our working hypothesis is that release of the amino acid substrate is facilitated by deprotonation of a key glutamate residue (E115), located at the bottom of the binding pocket and which forms part of the intracellular gate, allowing the protein to transition from an inward-occluded to an inward-open conformation. During unbiased molecular dynamics, we observed a transition from the inward-occluded state captured in the crystal structure, to a much more open state, which we consider likely to be representative of the inward-open substrate release state. To explore this and the role of protons in these transitions, we have used umbrella sampling to demonstrate that the transition from inward-occluded to inward-open is more energetically favourable when E115 is deprotonated. That E115 is likely to be protonated in the inward-occluded state and deprotonated in the inward-open state is further confirmed via the use of absolute binding free energies. Finally, we also show, via the use of absolute binding free energy calculations, that the affinity of the protein for alanine is very similar regardless of either the state or the protonation of E115, presumably reflecting key interactions deep within the binding cavity. Together, our results give a detailed picture of the role of protons in driving one of the major transitions in this transporter.

**Significance Statement:** For transporter proteins that utilize the proton gradient to drive the uptake of solutes, the precise mechanistic details of proton-coupling remain poorly understood. Structures can only infer the position of protons. All-atom molecular dynamics simulations however, are the ideal complementary tool. Here, we report extensive MD simulations on GkApcT, a proton-coupled transporter. We observe a spontaneous transition from the crystallographically derived inward-occluded state, to an inward-open state, which we then characterise with umbrella sampling and absolute binding free energy calculations. The results suggest that a conserved glutamate is protonated in the inward-occluded state and subsequent deprotonation of this glutamate allows the transporter to move into the inward-open state, thus facilitating substrate release into the cell.

## Introduction

The intracellular concentrations of amino acids are regulated by various transporters that utilize different mechanisms. Transport can in some cases be coupled to another electrochemical gradient, such as sodium (1) or protons (2), which are used to drive the concentrative uptake of the amino acid into the cell. Proton coupled amino acid transport systems, which include the SLC36 family of PAT transporters, are also involved in mediating the export of amino acids from the lysosome in eukaryotic cells, and in certain instances, have been shown to regulate cellular metabolism through interactions with the mechanistic target of rapamycin (mTORC) kinase signaling complex (3, 4). Some transporters can also act as facilitators, whereby the substrate is exchanged for its metabolized product (5) or another amino acid (6). This exchange activity can also be regulated by protons (pH) where the transporter is only active in acidic pH (7). Although there have been many hypotheses put forward concerning the overall conformational cycle of secondary active transport proteins (8) the precise details of how the structure of these membrane proteins and their associated transport activity is regulated by protons remains incomplete.

A proton-coupled amino acid transporter with the LeuT fold from *Geobacillus kaustophilus* (GkApcT) was recently solved in an inward-occluded state (9). GkApcT is distantly related to both the SLC7 family of cationic amino acid transporters and the SLC36 family of proton coupled amino acid transporters. As such, it serves as a useful model system to understand how protons can be used to drive the transport of amino acids across cellular membranes. Similar to other members of the Amino Acid Polyamine Cation (APC) transporter superfamily, the structural fold of GkApcT is comprised of an inverted repeat of two five-helix bundles, with an additional two extra helices in the C terminal (**Fig. 1A**). A common view is that transporters that share this structural fold share similar transport mechanisms such as those outlined in **Fig. 1B**. The binding of the substrate to the outward-open state leads to the closure of the extracellular gate giving rise to the outward-occluded state. The transporter then enters a state where the extracellular gate is firmly closed and the intracellular gate is primed to open. This state is termed the inward-occluded state and is followed by the opening of the intracellular gate and the release of the substrate, whereby the transporter is in the inward-open state and the intracellular gate is fully open.

**Figure 1.**
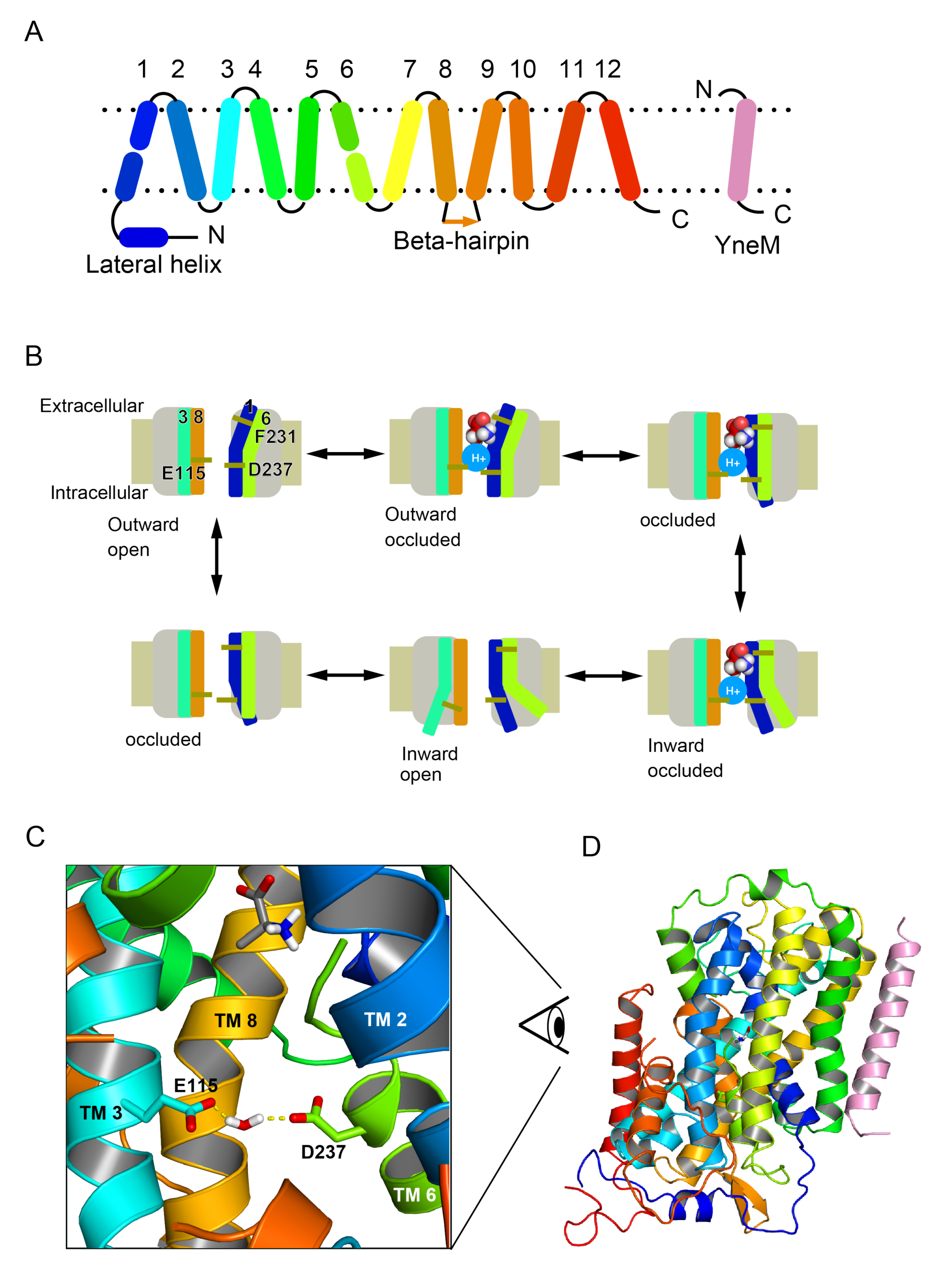
Structure and mechanism of GkApcT. (**A)** Schematic diagram of the GkApcT fold, where the first five helices are an inverted repeat of the next five helices, similar to LeuT. The YneM helix (Pink) is an additional subunit present in the crystal structure and included in the simulations presented here. (**B)** Mechanism of proton coupled transport cycle. 1. Apo outward-open state; 2. outward-open state with substrate and proton bound; 3. occluded state 4. Inward-open state with substrate and proton ready to leave; 4. Apo inward-open state; 6. Occluded state. (**C)** GkApcT in inward-occluded state in complex with alanine. The whole transporter is shown on the right and a magnified view of the binding pocket is shown on the left. The substrate alanine is drawn as grey sticks. The two key residues gating the intracellular gate are E115 from TM 3 (aqua stick) and D237 from TM6 (green). The water molecule is not directly observed in the crystal but appears in molecule dynamics simulations.

The crystal structure of GkApcT suggested that a glutamate residue (E115) within transmembrane helix (TM) 3, located at the intracellular side of the binding pocket, might be the key to the opening of the transporter towards the interior of the cell and the ejection of substrate. The intracellular gate, which sits underneath the amino acid binding pocket also contains an aspartate residue, located on TM 6 (D237), which appears to stabilise the closed state of the intracellular gate by forming an interaction with E115 through a bridging water molecule (**Fig. 1C**).

The role of protons in the transport mechanism of several transporters appears to direct the formation and breaking of conserved salt bridge interactions between the extracellular and in intracellular gating helices (10, 11). We previously hypothesised that the important location of E155 and D237 in stabilising the closed the state of the intracellular gate suggested these two side chains might play a similar role in GkApcT. The close position of E115 and D237 in combination with a calculated pKa value of 8.22 for E115, suggested this side chain was protonated in the crystal structure. Deprotonation on the intracellular side of the membrane should result in the repulsion of E115 from D237, driving the opening of the intracellular gate and release of the bound amino acid.

To explore this further and evaluate this hypothesis, we have used appropriate computational methods (12) to examine the protonation of E115 and the influence this has on the free energy landscape of the inward-occluded to inward-open transition. In unbiased MD simulations with E115 deprotonated, we were able to observe a spontaneous transition from the inward-occluded state to a new conformation state that is consistent with the properties expected for an inward-open state, including a clear exit path for the bound amino acid. We used this as the basis for potential of mean force and absolute binding free energies (ABFE) to characterize the free energy landscape of the inward-occluded to inward-open transition.

Our results support a mechanism whereby deprotonation of E115 is necessary and sufficient to drive the transition to the inward-open state, thus enabling release of the amino acid into the interior of the cell. The likely state of E115 protonation in either the inward-occluded or inward-open is further confirmed by ABFE and also suggests that the ligand does not strongly influence this preference. Finally, ABFE of alanine in the binding pocket in various states suggest that the affinity of the protein for alanine is not substantially influenced by the protonation states. Our results thus provide a detailed picture of the energetic landscape for the transition from inward-occluded to inward-open state in GkApct, and a framework for understanding related mechanisms within the SLC36 family of proton coupled transporters in mammalian cells.

## Methods

### System Preparation

The structure of the M321S GkApcT constructs (6F34) was used (13). The binding pose of alanine to M321S mutant was generated by truncating the arginine. All organic molecules except cholesterol were removed and the missing residues were modelled using modeller (14). A thousand models were generated and the best one is selected as the model with the best QMEANDisCo score (15). The missing residues were modelled using modeller and all organic molecules except cholesterol were removed. The crystal water bridging the C-terminus of the ligand and the protein was preserved, while all other crystal water was removed.

### Bilayer Setup

The GkApcT, YneM helix and cholesterol obtained from X-ray crystallography were converted to a coarse-grained (CG) representation. The conversion was performed using the MARTINI v2.2 coarse-grain forcefield (16), where 4 atomistic particles were represented as one “bead”. The self-assembly process was initiated by adding 500 1-palmitoyl-2-oleoylphosphatidylcholine (POPC) lipid molecules to the simulation box. Coarse-grained simulation was then performed for 25 ns in order for the lipids to form bilayer around the protein.

The POPC bilayer was converted to a mixture of POPE and POPG in a 3:1 ratio to match the experimental conditions. A box with protein in its centre and was twice as large as the bounding box of protein in all x, y, z dimensions was sculpted from the CG box. All the lipids outside this box were deleted. A 1 μs equilibration run was performed to allow POPE and POPG to equilibrate and the final frame was converted to atomistic resolution using the Backwards tool from the MARTINI toolkit (Wassenaar, Pluhackova et al. 2014).

The crystallographic coordinates of the GkApcT, YneM helix, cholesterol and substrate arginine or alanine were superimposed to the backward mapped POPE/POPG bilayer to replace their course-grain counterpart. The steric clash between the POPE/POPG lipid and the protein were resolved using the InflateGro methodology (17). The resulting bilayer system was solvated with TIP3P water (18) in a cubic box with periodic boundary conditions and neutralised with NaCl to reach a final ionic concentration of 150 mM. The resulting atomistic system was energy minimised using the steepest descents method and allowed to equilibrate for 25 ns in NPT ensemble with position restraint applied to GkApcT, YneM helix and cholesterol.

### All-atom Molecular Dynamics

Two replicates of 500 ns simulation were performed with E115 protonated and with E115 deprotonated and with alanine and arginine present. The AMBER ff99SB-ILDN forcefield (19) was used to describe protein and the Slipids forcefield (20) was used for the lipid molecules. The topology files for zwitterion arginine and alanine were obtained from AMBER parameter database (21). Protonation states of E115 were assigned using the pdb2gmx tool from GROMACS.

The timestep was 2 fs. The LINCS algorithm (22) was used to constrain H-bond lengths and the long-range electrostatic interactions were calculated with the PME method. The system was heated in NVT ensemble with V-rescale thermostat for 200 ps to reach a final temperature of 310K. The system was then equilibrated in NPT assemble for 5 ns with nose-hoover thermostat at 310K and the Berendsen barostat (23) to keep pressure at 1 bar. During the equilibration phase, all heavy atoms were position restrained at 1000kJ/mol. The production runs were performed under NPT ensemble and the temperature was kept at 310 K with nose-hoover thermostat (24) at 310K and the pressure at 1 bar with the Parrinello-Rahman barostat (25).

### Parameterization of Protonated Glutamate

As noted by Plamen Dobrev et al. (26), the rotation potential of the carboxyl H atom in Amber99SB-ILDN force field favours the syn orientation compared with the anti. Indeed, our preliminary simulations with default parameters confirmed this observation, which is also inconsistent with the potential surface obtained from QM calculations. Thus, we have reparameterised the C-C-O-H dihedral such that the potential energy in TIP3P matches calculations performed at the HF/6-31G(d) level of treatment with a PCM water solvent (using Gaussian).

Due to the instantaneous fluctuation of the water molecules, the potential energy of the system is taken as the average potential of the system across 100 ns simulation. During the simulation, a glutamate residue with N-terminus capped with acetyl group and C-terminus capped with N-methyl group was solvated with TIP3P water in a tricyclic box with edge length 4 nm. A sodium ion was also added to the box to neutralized the system. To minimize interaction between glutamate residue and the sodium ion, the sodium ion was positionally restrained to the corner of the box while the Cα of glutamate is restrained to the center of the box. An initial dihedral scan composed of 72 simulations with 5 degree steps, each with the C-C-O-H dihedral restrained at 3000 kJ/mol, were performed with the C-C-O-H dihedral potential set to zero. The resulting potential was fitted with a three-term Ryckaert-Belleman potential to match same dihedral potential surface obtained in Gaussian. The dihedral scan was computed again with the new fitted Ryckaert-Belleman potential to check that the final result is consistent with the QM dihedral potential surface.

### Umbrella Sampling Simulations and Potential of Mean Force Calculations

The collective variables for umbrella sampling were defined as the distance between E115 and D237. The coordinate of E115 was defined as the geometric centre of the alpha carbons of residue number 113 to 117. The coordinate of D237 was defined as the geometric centre of the alpha carbons of residue number 235 to 239. A total of 25 windows were created from 13 Å to 19 Å with a step of 0.25 Å. The initial frames for umbrella sampling were extracted from the unbiased simulation of GkApcT (M321S mutant) in complex with alanine with E115 deprotonated. The frame which was closest to the specified CV for each window was chosen as the initial frame for that window.

The umbrella sampling was performed in GROMACS 5.1.2 patched with Plumed 2.3. The bias applied to the collective variables was 1000kJ/mol. The PMF profiles were generated using WHAM (Grossfield, Alan). The free energy of conformational changed is defined as the difference between the inward-occluded state and the inward-open state. The inward-occluded state is defined as the point of lowest energy in the free energy profile with E115 protonated, whereas the inward-open state is defined as the same lowest energy point when E115 is deprotonated. The error estimation is done in block analysis fashion, whereas WHAM analysis is performed with the last 50%, 40%, 30%, 20% and 10% of the data. The error of the result is estimated as the standard deviation of these five values.

### Alanine Absolute Binding Free Energy Calculations

Calculations were performed in GROMACS 2018 and followed a standard alchemical free energy cycle as previously described by us (27, 28). The charges were annihilated with Δλ= 0.25, whereas 15 non-uniformly distributed λ values (0.05, 0.1, 0.2, 0.3, 0.4, 0.5, 0.6, 0.65, 0.7, 0.75, 0.8, 0.85, 0.9, 0.95, 1.0) were used for the decoupling of van der Waals interactions. Similarly, to restrain the substrate to the protein, 10 non-uniformly distributed λ values (0.01, 0.025, 0.05, 0.075, 0.1, 0.2, 0.35, 0.5, 0.75, 1.0) were used. Thus, a total of 36 windows for the substrate and protein complex simulation and ligand simulation were employed. A timestep of 2 fs was used for the restraint free energy and the free energy of charge annihilation. A 1 fs timestep was used for the decoupling of the ligand and 0.5 fs timestep was used for systems near the end of the decoupling. The conformations of the protein were restrained by using a harmonic restraint of 1000 kj/mol on the collective variable as described above for the umbrella sampling. The value of the collective variable for each conformation were defined as the distance of the lowest point in the PMF. After equilibration, dH/dl data were collected from 30 ns of production runs. The relative position and orientation of the substrate to the protein were restrained using one distance, two angles and three dihedrals and the restraint free energy was evaluated analytically (29). The collected dh/dl were processed using alchemical analysis (30).

### Protonation Free Energy Calculations

Calculations were carried out in GROMACS 2018.1 similar to the absolute binding free energy calculations, where the protonated glutamate was alchemically transformed to deprotonated glutamate. The alchemical charge transformation is done from protonated glutamate to deprotonated glutamate with Δλ = 0.1. Five lambda windows (Δλ = 0.25) were also used to transform the bond length, angle and mass all at the same time. Thus, a total of 16 windows were run for 30 ns and the result was analysed with alchemical analysis. To balance out the finite size problem introduced by the non-neutral box at the end state, a chloride ion was alchemically transformed at the same time to keep the box neutral. To prevent interaction between the glutamate and chloride ion, the chloride ion is position-restrained to the corner of the box ensuring an 80 Å separation between the two alchemically transform objects.

The free energy of annihilation of the charge of the chloride ion were deducted from the final result. The charge annihilation free energy was initially obtained from simulating a chloride ion in a cubic box of water. The charge of the chloride ion was annihilated with Δλ = 0.1. The simulation setup is the same as all other free energy calculations. The Rocklin correction (31) was applied to correct for the size-dependent finite-size error. Different box sizes with edge length of ranging from 30 Å to 100 Å with a step of 10 Å were tested to ensure the final corrected charge annihilation free energy is size independent. For these 9 box sizes, three replicates were performed and the final chloride ion charge annihilation free energy is taken as the average of the 9 × 3 trials.

## Results

### Protonation of E115 stabilizes the close state of the transporter

GkApcT functions to couple amino acid uptake to the inwardly directed proton electrochemical gradient (9). The crystal structure was captured in a so-called inward open state, whereby the extracellular gate was tightly sealed, and the intracellular gate partially open but closed enough to occlude the bound amino acid within the central binding site. The transition from the inward-occluded state to the fully inward-open state was postulated to involve an increase in the separation between TMs 3 and 6 (**Fig. 1B**). Given the location of E115 on TM3, and its unusual pKa of 8.2 compared to 4.38 for D237 (calculated by propKa), we postulated the conformation of the intracellular gate is likely dependent on the protonation state of this side chain (9).

To investigate this in the first instance we performed a series of unbiased 500 ns MD simulations of GkApcT in a POPE/POPG (3:1 ratio) lipid bilayer (**Fig. 2**) with E115 in different protonation states and in the presence and absence of an amino acid (either alanine or arginine - **see Methods**). During one of the simulations with E115 deprotonated, we observed a significant expansion of the binding cavity (**Fig. 2B**) along with an increase in the distance between TM3 and TM6 (**Fig. 2C**) at the intracellular end of the helices, resulting in a state that would be consistent with an inward open conformation. In contrast, this movement was not observed when E115 remained protonated.

**Figure 2.**
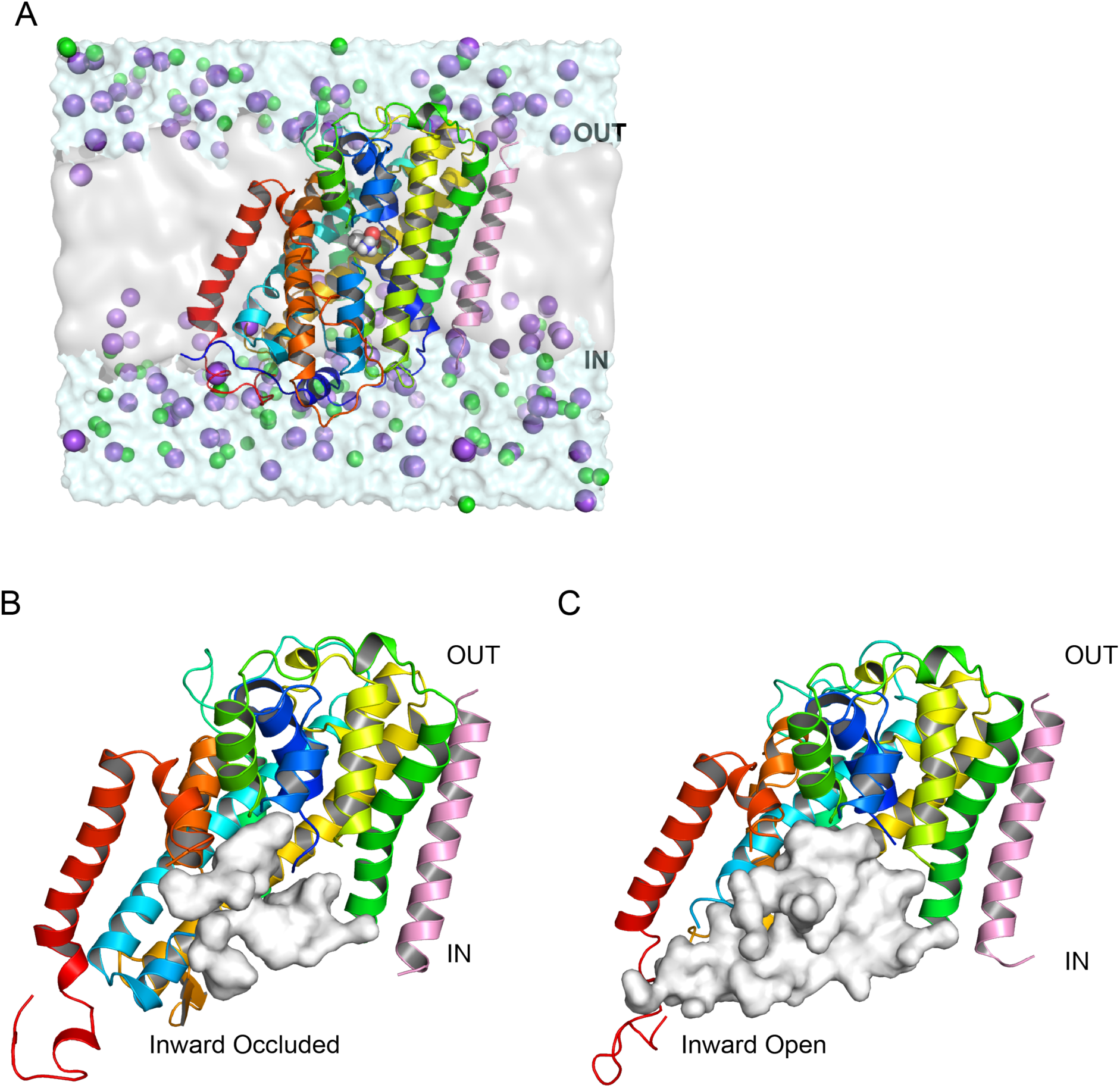
Unbiased molecular dynamics simulations of GkApcT in the lipid bilayer. **(A)** Snapshot of the simulation system - GkApcT embedded in POPE/POPG bilayer with alanine sitting in the binding pocket. **(B)** GkApcT in inward-occluded conformation and (**C**) inward-open conformation (right). The conformational change from inward-occluded to inward-open results in an expansion of the binding cavity as represented by the white surface. In the inward-open conformation, the binding cavity extends to the bulk of intracellular space giving rise to a potential substrate exit pathway

### Transition to an inward open state following deprotonation of E115

To characterize the new conformational state and to establish whether indeed it could be considered a possible inward-open state, we first pulled the alanine out of the binding pocket using steered MD with a moving restraint that increased the distance between the binding pocket, defined as the centre of mass of the Cα of residues I40, T43, G44, F231 and I234 (i.e. the residues that make hydrogen-bonds to the bound alanine in the binding site) and the Cα atom of the alanine. **Fig. 3A** shows the profile of the work done for five such pulling simulations as a function of this vector. The plateaus (at 60 ns and beyond in all simulations) correspond to the point in time when there are no hydrogen-bond interactions between the alanine and the residues of the binding site, and the ligand exhibits random motion within the cavity. Although these profiles are far from converged, we were ultimately focused on whether there were further impediments to access to the bulk solution beyond that of the binding site. These simple pulling simulations show there are none, and the amino acid remains bound only through the interactions with the binding site residues.

**Figure 3.**
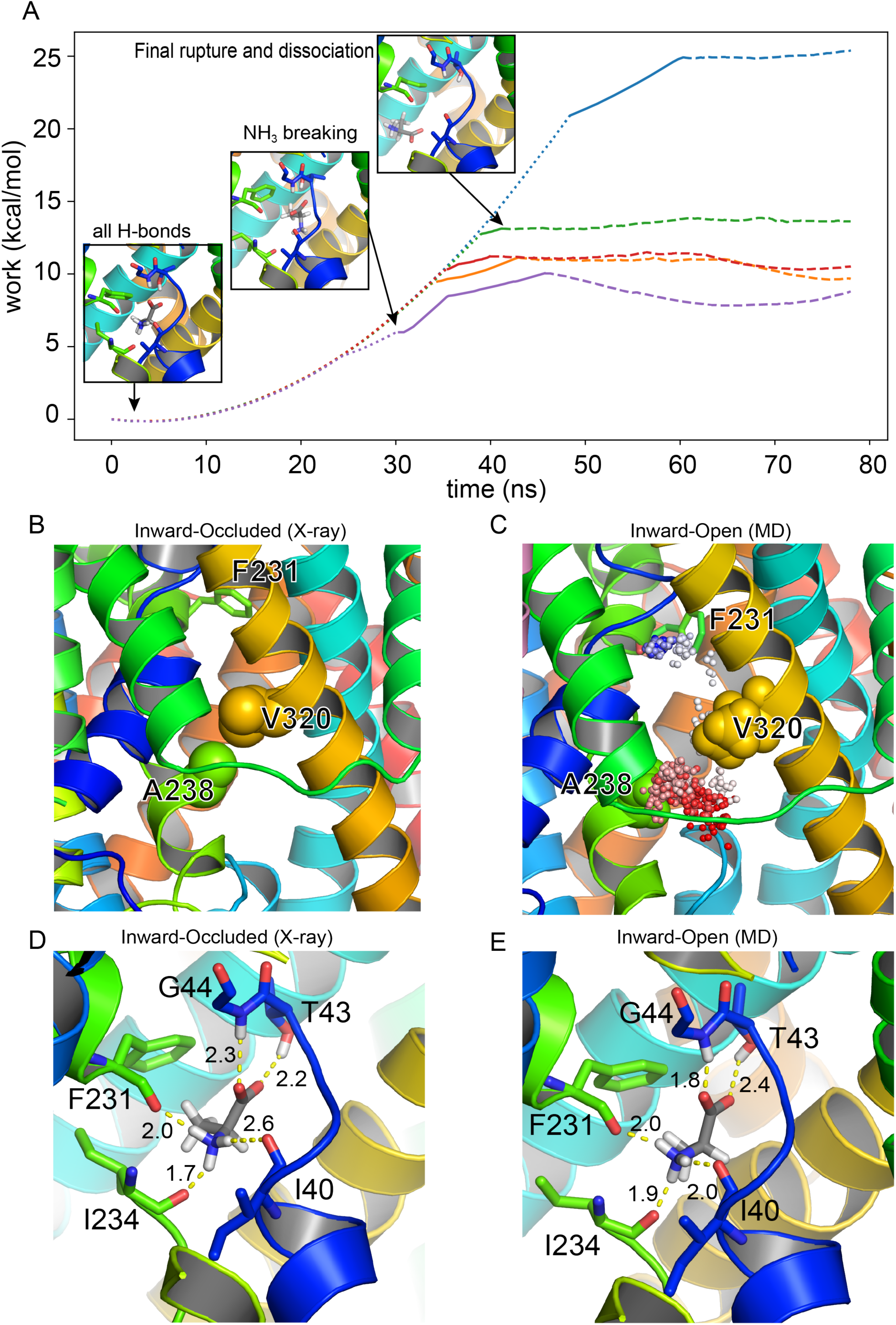
**(A)** Work performed as function of time in the pulling simulation. The dotted section of the line represents the state where alanine is fully coordinated in the binding pocket. The solid section is where the interaction between the amine group of alanine and the protein is broken while the interaction between the carboxyl group and protein is still intact. The dashed line represents the state where alanine is fully dissociated from the binding pocket. Snapshots give an indication of the nature of the binding event at the various transition points. The plateaus reflect the alanine diffusing randomly within the cavity away from the binding site. (**B**) shows the gate in the inward-occluded state formed by A238 and V320 drawn as spheres (not no hydrogens are present in the crystal structure). (**C**). The path (as shown as the coloured dots from red to blue as a function of time) of the alanine in an unbiased simulation that show re-binding to the binding site (indicated by the F231 residue) during the inward-open state. (**D**, **E**) show a comparison of alanine binding to the inward-occluded and inward-open states respectively.

To explore this further, we took one snapshot from the end of one of the pulling simulations, which we considered was fully dissociated from the binding pocket, and performed free molecular dynamics from this start point. After 180 ns of simulation time, we observed the spontaneous re-binding of alanine to the binding pocket (**SI movie SI-M1**). The main barrier to the exit of alanine from the inward-occluded state is a hydrophobic gate formed by A238 on TM6 and V328 on TM8. In the crystal structure, these act (**Fig. 3B**) to occlude access to the bulk solution. In the inward-open state however, the separation of these helices creates a large enough space to allow alanine to move quite freely (the closest separation is between atoms of A238 and V328 is about 6.5 Å) as is shown in **Fig. 3C** alongside a timeline trace of the path of alanine as it moves back into the binding pocket. Closer inspection of the binding event revealed that the binding mode is practically identical to that observed in the crystal structure (**Fig. 3D and E**).

Thus, this observation and the steered MD profiles confirm that there is no steric barrier to ligand egress from the binding site and thus the final state sampled in our simulations is likely to be reflective of a genuine inward-open state.

### Potential of Mean Force Calculations identify E115 as an important part of the proton coupling mechanism

Satisfied that the opening we observed represented a plausible inward-open state, we decided to use umbrella sampling (see **Methods**) on the various systems (E115 protonated/deprotonated, alanine present/absent) to further map out the energy landscape of this conformational change. We used snapshots from the transition observed in the unbiased simulations as the starting points and constructed a collective variable defined as the distance between the centre of mass of the Cα atoms of residues 113 to 117 on TM3 and the centre of mass of the Cα atoms of residues 235 to 239 on TM6. PMF profiles were calculated for this collective variable (see **Methods**). **Fig. 4** and **Table 1** summarize the calculations for all systems.

**Table 1.** Summary of free energy differences in kcals/mol between inward-open and inward-occluded states.

| | $\Delta G$ [E115] | $\Delta G$ [E115H] | $\Delta\Delta G$ |
| --- | --- | --- | --- |
| Apo | $-3.13 \pm 0.1$ | $1.33 \pm 0.2$ | $-4.46 \pm 0.2$ |
| Alanine | $-2.04 \pm 0.1$ | $2.64 \pm 0.1$ | $-4.68 \pm 0.1$ |

$\Delta G$ is the free energy difference between inward-open and inward-occluded state ( $G_{\text{inward-open}} - G_{\text{inward-occluded}}$ ). $\Delta\Delta G$ is the free energy difference of the $\Delta G$ in different protonation states. This is calculated as the difference between the free energy of the conformational change when E115 is deprotonated ( $\Delta G[\text{E115}]$ ) and the free energy of the same conformational change when E115 is protonated ( $\Delta G[\text{E115H}]$ ). Errors are estimated by block averaging (see methods).

**Figure 4.**
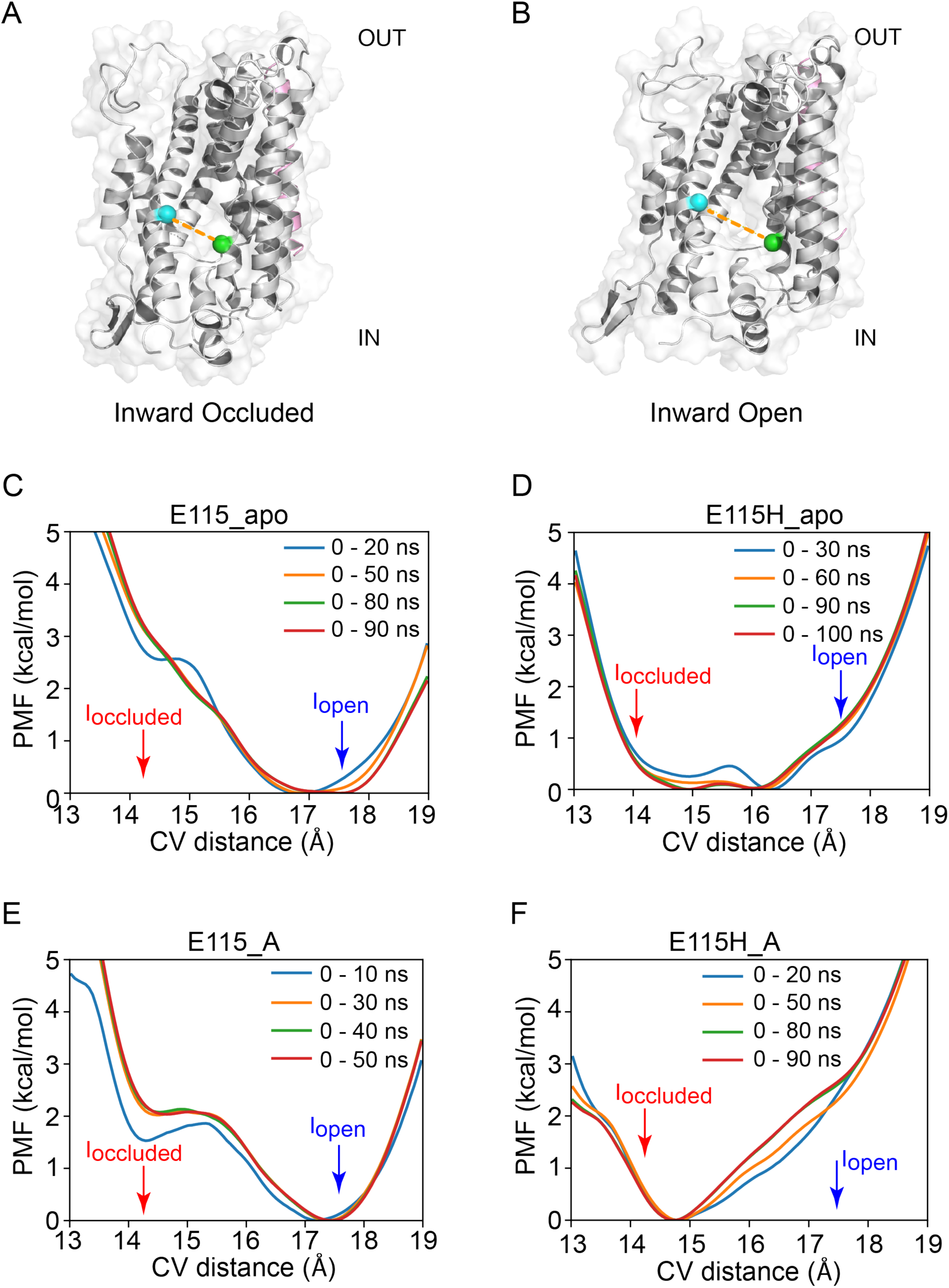
**(A and B)** The space between two green spheres represents E115 and D237. The distance between the two spheres increases as GkApcT transitions from inward-occluded (**A**) to inward-open (**B**). This distance formed the basis of the collective variable used in later umbrella sampling calculations. (**C-E**) Potential of mean force calculation using umbrella sampling. The x-axis is the collective variable, which is the distance between the centre of mass of the Cα atoms of residues 113 to 117 and the centre of mass of the Cα atoms of residues 235 to 237. The relative locations of the inward-occluded and inward-open state are annotated by red and blue arrows respectively. PMF profiles are shown for (**A**) Apo with deprotonated E115, (**B**) Apo with protonated E115, (**C**) Deprotonated with alanine in the binding site, (**D**) Protonated E115 with alanine bound, Different lines show how the profiles change until convergence in each case, defined as no discernible difference after further sampling. Bootstrap errors are smaller than the width of the lines.

The PMF profiles (**Fig. 4**) show that energetic cost of moving from inward-occluded to inward-open is dependent on the protonation state of E115. The conformational change from inward-occluded to inward-open is energetically favourable when E115 is deprotonated, where the free energy differences (ΔG = G_inward-open_ - G_inward-occluded_) were negative (**Fig. 4 A and C** and **Table 1**). Deprotonation of E115 favours the inward-open state (**Fig. 4A, C**) by ∼ 3 kcals/mol in the apo state and 2.2 kcals/mol when alanine is in the binding site. In comparison, the PMF exhibits a much larger, flatter well when E115 is protonated in the apo state (**Fig. 3B**) and although the inward-occluded and inward-open states are not located at the bottom of the well, the energies associated with these positions is slightly larger than 1 kcal/mol. When alanine is present, the PMF exhibits a sharper well profile, with the minimum located very close to the inward-occluded state. The energy associated with the inward-open state is of the order of 2.6 kcal/mol, consistent with the expectation that ligands would be expected to stabilize the inward-occluded state, but not the inward-open state.

### Absolute Binding Free Energies confirm a role for protonation of E115 in stabilising the inward open state

Although the PMF calculations gave results consistent with current hypotheses on proton-amino acid transporters, we sought to independently verify the energetics via the use of ABFE. The standard thermodynamic cycle approach was used (see **Methods**, **Tables 2 and 3**). We first use ABFE to calculate the free energy associated with protonating E115 (**Table 2**). The results suggest that a protonated E115 residue helps to stabilize the inward occluded state, completely consistent with the results obtained from the PMF calculations.

**Table 2.** Summary of free energy to *protonate* the E115 sidechain. All energies in kcal/mol.

|  | Inward-occluded | Inward-open |
| --- | --- | --- |
| Apo | $-3.62 \pm 0.2$ | $+2.72 \pm 0.1$ |
| Alanine bound | $-6.01 \pm 0.2$ | $+7.01 \pm 0.2$ |

Errors are estimated by MBAR under error propagation (see methods).

**Table 3.** Absolute free energies for alanine. All energies in kcal/mol ± standard deviation.

|  | Alanine in water | Alanine in E115_Inward-occluded | Alanine in E115_Inward-open | Alanine in E115H_Inward-occluded | Alanine in E115H_Inward-open |
| --- | --- | --- | --- | --- | --- |
| $\Delta G$ | $-47.73 \pm 0.01$ | $-5.89 \pm 0.04$ | $-5.55 \pm 0.07$ | $-5.67 \pm 0.08$ | $-5.27 \pm 0.14$ |

*\*Errors are estimated by MBAR under error propagation (see methods).*

We then used ABFE to compute the free energies of alanine binding to the different conformational states and different protonation states. Those results are summarized in **Table 3**. Interesting, it can be seen that protonation of E115 does not change the affinity of alanine for the inward-occluded state. The affinity of alanine for the inward-open state is slightly reduced, but only by 0.5 kcal/mol. Too small to indicate any significant difference to how the amino acid is recognised within the transporter. The largest change in affinity is in the inward-open state with E115 protonated, which gives an affinity for alanine of −4.6 kcal, which is only 1 kcal/mol difference.

## Discussion

Molecular dynamics (MD) has emerged as a powerful tool to understand the role of ion binding to several members of the APC superfamily, including the sodium coupled transporter LeuT (32, 33). However, GkApcT is coupled to protons rather than sodium and the question of the how the same protein fold has evolved to couple to different ion gradients is ongoing within the transporter community (34). The observation that members of the SLC36 family of proton coupled amino acid transporters regulate the mTORC complex and shuttle between the plasma membrane and lysosomal membranes, which both contain a pH gradient across them, further underscore the need to understand the role of protons in regulating transporter function within the cell.

MD simulation is an ideal tool to study this kind of problem, but one must be careful, especially when trying to interpret short timescale events in the simulations and their relevance to longer time-scale observables. Indeed, the implicit assumption of many MD studies that local conformational changes observed in short simulations can provide insight into global motions has recently been called into question (35). It is clear that very long simulation times are often required, as recently demonstrated for the semiSWEET transporter (36) or the GluT1 transporter (37). Indeed, to study the complete conformation cycle, it would seem likely that the approach of Markov State Modelling (MSM) might be the most appropriate as recently demonstrated with PepT_so_ transporter (38). In this work however, we were interested in fully characterizing only one part of the full transition pathway, facilitating the use of shorter timescales that enabled the application of more rigorous methods.

We first examined whether the deprotonation of E115 facilitates opening of intracellular gate in GkApcT. The PMF calculations show that the free energy differences of the conformational change were more negative, i.e. more favourable, when E115 was deprotonated, consistent with our earlier hypothesis (9). However, our results also show that this conformational change is necessary in order for the substrate to exit (or enter) the binding pocket from the interior of the cell in unbiased simulations. Although PMF calculations have been reported to obtain a reasonable estimate of unbinding (for example xylopyranose from XylE (39)) we could not rule out alternative dissociation pathways and hence elected to use ABFE to support our PMF calculations. Using ABFE, we have shown that protonation of E115 stabilizes the inward-occluded state and that this is the case regardless of the presence of ligand in the binding site. The ABFE calculations for alanine affinity were initially surprising to us, in that they did not reveal weaker binding to the inward-occluded state as one might perhaps expect. However, when one considers that the key interactions between the protein and ligand are deep in the cavity and do not appear to change much between the inward-occluded and inward-open states (**Fig. 3 D, E**) then the lack of difference is less surprising. It was recently suggested for LeuT, via the use of single-molecule fluorescence resonance energy transfer (smFRET), that partially open intermediates were associated with transport activity (40). We would suggest a note of caution here, as although our inward-open state appears seductive, we do not know whether a more open (or different) inward-open state with more significant changes in the binding might be possible. We should also point out that it might be tempting to combine all the calculations to generate a full thermodynamic cycle that links all of the possible states. However, the calculation of protonation states with a fixed charged model as we have performed here, is well known to be problematic (41, 42) and more work is clearly needed in this area.

Another question that remains at this point is where the proton moves to after it leaves E115. That is beyond the scope of the current work and dealing with proton-hopping in complex systems is far from trivial although there is promising work in this direction (43). However, it is likely to involve water and indeed the role of water in driving conformational transitions is made very apparent here. Water penetration into the inward-occluded state will facilitate the deprotonation step and accelerate the probability of moving into the inward-open state. A similar role has been proposed on the basis of MD simulations for the DAT (44), SERT (45), the XylE *(46)* and LeuT transporters (47, 48) whereby water penetration leads to release of the Na2-bound Na^+^ which in turn leads to destablization of the substrate. It would be interesting to examine this further with constant pH simulations as has been demonstrated for the sodium-proton antiporter (49, 50).

## Conclusions

We have provided mechanistic insight into the role of proton-coupled transport in GkApcT. Specifically, we have developed a model for how the proton and substrate are released from the inward-occluded state observed previously in GkApcT. Our results provide further evidence that deprotonation of E115 is associated with the opening of the intracellular gate as well as the transition from inward-occluded to inward-open state. This conformational change opens up the passage from the binding pocket to the intracellular space allowing the release of the substrate. Importantly, E115 is conserved in the mammalian SLC36 homologues, suggested this residue is likely to form part of a more conserved mechanism for proton coupling within the APC superfamily.

## Author Contributions

PCB and SN designed the project. ZW performed the calculations. IA advised on PMF and FEP interpretation. IA, ZW, SN and PCB analysed the data. ZW, SN and PCB wrote the manuscript.

## Acknowledgements

SN & ZW thank the Wellcome Trust (203741/Z/16/Z & 102890/Z/13/Z) for support. This project made use of time on JADE (EP/P020275/1) and ARCHER granted via the UK High-End Computing Consortium for Biomolecular Simulation, HECBioSim (http://www.hecbiosim.ac.uk), supported by EPSRC (EP/L000253/1).

